# ssDNA is not superior to dsDNA as long HDR donors for CRISPR-mediated endogenous gene tagging in human diploid cells

**DOI:** 10.1101/2022.06.01.494308

**Authors:** Akira Mabuchi, Shoji Hata, Mariya Genova, Chiharu Tei, Kei K Ito, Masayasu Hirota, Takuma Komori, Masamitsu Fukuyama, Takumi Chinen, Atsushi Toyoda, Daiju Kitagawa

## Abstract

Recent advances in CRISPR technology have enabled us to perform gene knock-in in various species and cell lines. CRISPR-mediated knock-in requires donor DNA which serves as a template for homology-directed repair (HDR). For knock-in of short sequences or base substitutions, ssDNA donors are frequently used among various other forms of HDR donors, such as linear dsDNA. However, for insertion of long transgenes such as fluorescent reporters in human cells, the optimal type of HDR donors remains unclear. In this study, we established a simple and efficient CRISPR-mediated knock-in method for long transgenes using linear dsDNA and ssDNA donors, and systematically compared the performance of these two donors for endogenous gene tagging in human non-transformed diploid cells. Quantification using flow cytometry revealed higher efficiency of fluorescent tagging with dsDNA donors than with ssDNA. By analyzing knock-in outcomes using long-read amplicon sequencing and a classification framework, a variety of mis-integration events were detected regardless of the donor type. Importantly, the ratio of precise insertion was higher with dsDNA donors than with ssDNA. Moreover, in off-target integration analyses, dsDNA and ssDNA were comparably prone to non-homologous integration. These results indicate that ssDNA is not superior to dsDNA as long HDR donors for gene knock-in in human cells.

## Introduction

Gene knock-in is a crucial technique for studying gene function by introducing specific mutations or insertions at endogenous loci. Recent developments in genome editing technology using programmable site-specific nucleases, especially the CRISPR-Cas system, have made it possible to perform gene knock-in in a broader range of species and cell lines (Hsu et al., 2014). Cas9 and Cas12a nucleases, which are used for CRISPR-mediated genome editing, are targeted to specific genomic loci with short guide RNA to induce double-strand breaks (DSBs) (Gasiunas et al., 2012; Jinek et al., 2012; Zetsche et al., 2015). These DSBs can be repaired by two major pathways. The first is the re-ligation of the broken DNA ends through non-homologous end joining (NHEJ). This pathway is error-prone and often introduces insertions or deletions (indels), which can lead to gene knockout (Chang et al., 2017). The second pathway is homology-directed repair (HDR), in which DSBs are repaired precisely by using homologous DNA sequences as a repair template (Yeh et al., 2019). In HDR, exogenously introduced DNA can also serve as a repair template when the sequence contains so-called homology arms (HAs) - elements homologous to the region flanking the target site. Precise gene knock-in therefore requires exogenous donor DNA to utilize the HDR pathway.

The optimal type of DNA donors used as HDR templates for gene knock-in would depend on the length of the sequence to be inserted at the targeted site. For knock-in of short sequences or base substitutions such as point mutations, single-stranded oligodeoxynucleotides (ssODNs) are frequently used due to their high knock-in efficiency and ease of synthesis (Chen et al., 2011; Wu et al., 2013; Zhang et al., 2022). However, the optimal type of HDR donors for insertion of longer transgenes such as fluorescent reporters remains unclear. Various forms of donors, including plasmids, linear dsDNA produced by PCR, and ssDNA, are applicable for knock-in of long sequences. Although plasmids have been used as a conventional HDR donor, their preparation requires time-consuming cloning steps. Moreover, it has been reported that plasmids are less efficient than linear dsDNA or ssDNA for fluorescent tagging in human cell lines (Li et al., 2017; Paix et al., 2017).

Besides efficiency, the specificity and accuracy of the knock-in are also key factors that determine donor performance (Maggio and Gonçalves, 2015). It is known that exogenous DNA can be non-specifically inserted via non-HDR pathways into unintended locations of the genome, such as off-target cleavage sites introduced by Cas nuclease (Fueller et al., 2020; Tsai et al., 2015; Zelensky et al., 2017). Homology-independent donor integration can also occur at the target site of knock-in, which results in inaccurate insertion of the transgenes (Canaj et al., 2019; Renaud et al., 2016; Roberts et al., 2017). While both linear dsDNA and ssDNA donors can be inserted into the genome in a homology-independent manner, it has been reported that ssDNA donors are less prone to off-target integration than dsDNA (Chen et al., 2011; Li et al., 2017; Roth et al., 2018). In terms of accuracy, another report suggests that it depends on the cell type as to whether dsDNA or ssDNA donors would have a higher frequency of precise insertion via HDR at the target locus (Canaj et al., 2019). Thus, it remains controversial whether linear dsDNA or ssDNA templates are more suitable as HDR donors for insertion of long transgenes.

In this study, we compare the performance of dsDNA and ssDNA as long HDR donors for the endogenous tagging with fluorescent proteins in the hTERT-immortalized RPE1 cell line, which is one of the most widely used human non-transformed diploid cell lines. Quantitative analyses of the endogenous tagging in different genes show that ssDNA tends to have lower knock-in efficiency than dsDNA. It also turns out that ssDNA is not superior to dsDNA in terms of the specificity and accuracy of long transgene insertions. Taken together, our findings indicate that dsDNA is more suitable than ssDNA as long HDR donors for endogenous gene tagging with long sequences in human diploid cells.

## Results

### An optimized CRISPR knock-in method using long dsDNA donors for efficient tagging with fluorescent proteins in human diploid RPE1 cells

We first established a simple, long-dsDNA-based method for endogenous gene tagging with Cas12a or Cas9 by optimizing conventional approaches (Fig. 1a) (Fueller et al., 2020; Ghetti et al., 2021). Long dsDNA donors were amplified by a one-step PCR using a pair of primers containing 90 bases of HA sequences. To avoid plasmid construction, we synthesized the guide RNA (crRNA for Cas12a and sgRNA for Cas9) via *in vitro* transcription from PCR-assembled DNA templates. The guide RNA was mixed *in vitro* with recombinant Cas12a or Cas9 proteins to form ribonucleoprotein (RNP) complexes, which were then electroporated into RPE1 cells together with the long dsDNA donors.

**Fig.1:**
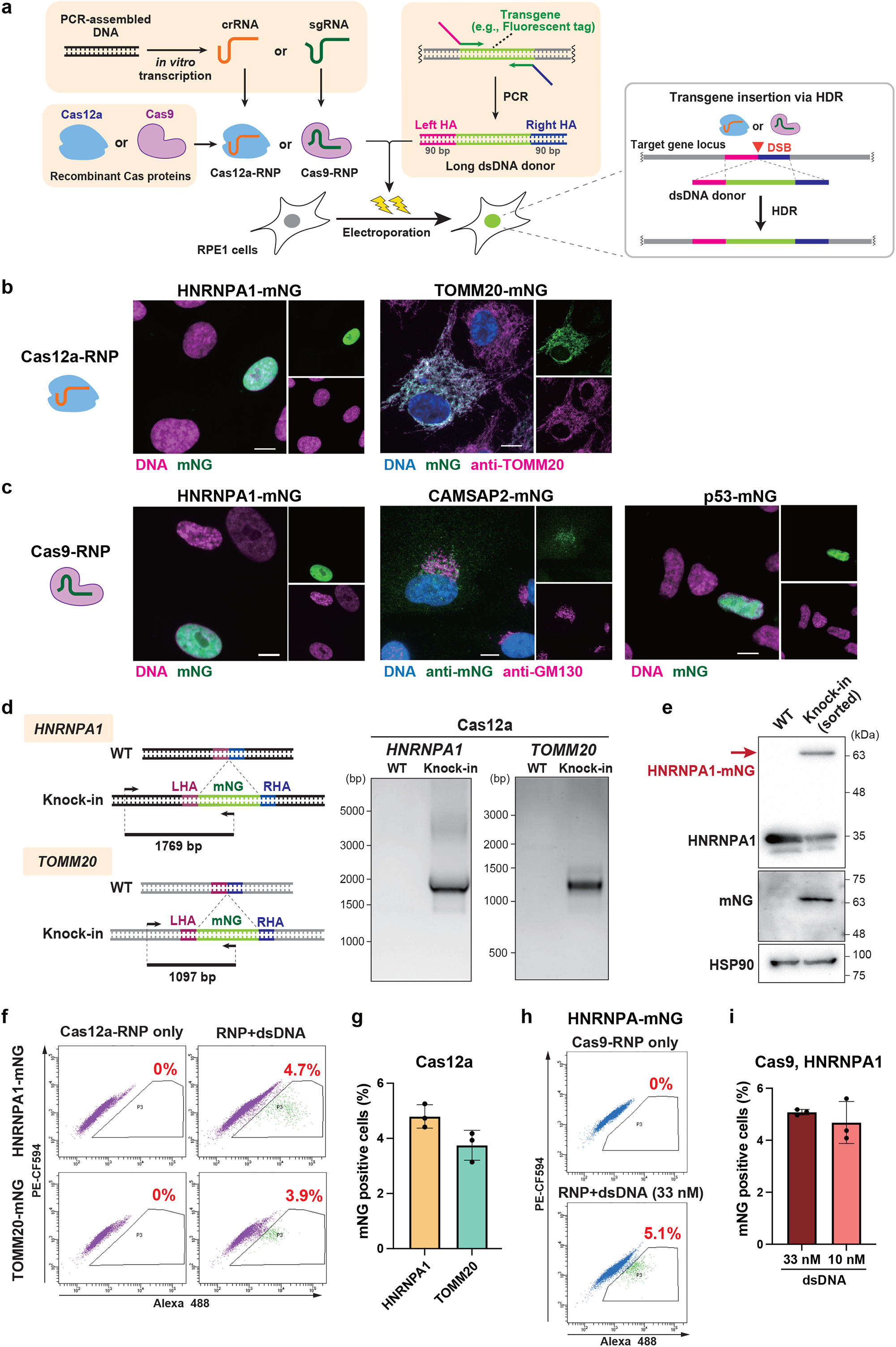
An optimized knock-in method using long dsDNA donors for efficient endogenous tagging with fluorescent proteins in human diploid RPE1 cells. **a**, Schematic overview of long-dsDNA-based endogenous gene tagging in human RPE1 cells. The long dsDNA donor is amplified by PCR using primers containing 90 bases of HAs. The guide RNA transcribed *in vitro* from PCR-assembled DNA is mixed with recombinant Cas12a or Cas9 proteins to form RNP complexes, which are electroporated with the dsDNA donor into RPE1 cells. The transgene is expected to be inserted via the HDR pathway into the target locus of the Cas-RNP using the dsDNA donor as a template. **b**, Representative images of cells with Cas12a-mediated endogenous mNG tagging of the indicated genes. Cells at 7-12 days after electroporation were fixed and analyzed. Scale bar: 10 µm. **c**, Representative images of cells with Cas9-mediated endogenous mNG tagging of the indicated genes. Cells at 12-17 days after electroporation were fixed and analyzed. Scale bar: 10 µm. **d**, Genomic PCR detecting the mNG insertion into the *HNRNPA1* or *TOMM20* locus with Cas12a-mediated knock-in. The primers were designed to amplify the 5’ junction of the mNG insertion for each gene. LHA: left HA, RHA: right HA. **e**, Western blotting confirming the fusion of mNG to HNRNPA1 via the Cas12a-mediated knock-in method. The knock-in cells were sorted by flow cytometry to collect mNG positive cells. HSP90 was used as a loading control. **f**, Flow cytometric analysis of Cas12a-mediated HNRNPA1-mNG and TOMM20-mNG knock-in cells. Cells at 8 days after electroporation were analyzed. Percentages of cells with mNG signal are shown in the plots. **g**, Quantification of percentages of mNG positive cells from (**f**). Data from three biological replicates are shown. >5,000 cells were analyzed for each sample of HNRNPA1 and TOMM20. Data are represented as mean ± S.D. **h**, Flow cytometric analysis of Cas9-mediated HNRNPA1-mNG knock-in cells. Cells at 5 days after electroporation were analyzed. Percentages of cells with mNG signal are shown in the plots. **i**, Quantification of percentages of mNG positive cells from (**h**). Two different concentrations of the dsDNA donor were analyzed. Data from three biological replicates are shown. 10,000 cells were analyzed for each sample. Data are represented as mean ± S.D.

To test whether this cloning-free method with the long dsDNA donors can be applied to efficient knock-in in RPE1 cells, we performed Cas12a-mediated endogenous tagging of the nuclear protein HNRNPA1 and the mitochondrial protein TOMM20 with the green fluorescent protein mNeonGreen (mNG). Fluorescence imaging revealed the expected localization of each mNG-fused endogenous protein (Fig. 1b). Similarly, Cas9-based mNG knock-in was carried out to target three different proteins (HNRNPA1, CAMSAP2, and p53). For each protein, a specific localization pattern of mNG was observed, indicating successful tagging of the targets (Fig. 1c). The Cas12a-mediated insertion of the mNG sequence into the target locus was confirmed by genomic PCR for the *HNRNPA1* and *TOMM20* genes (Fig. 1d). The specificity of the mNG tagging of HNRNPA1 was further confirmed by western blotting with antibodies against HNRNPA1 and mNG (Fig. 1e).

To evaluate the tagging efficiency of our approach, we conducted a flow cytometric analysis, which allows the detection of mNG positive cells in a high-throughput and quantitative manner. The analysis, performed on a cell population in which HNRNPA1 or TOMM20 was targeted for mNG tagging by Cas12a, revealed that the knock-in efficiency was 3 to 5% for each gene (Fig. 1f, g). A comparable level of mNG positive cells was detected using a similar strategy with Cas9 (Fig. 1h, i). Taken together, our optimized cloning-free knock-in method with long dsDNA donors enables efficient endogenous gene tagging in RPE1 cells.

### Production of long ssDNA donors with high purity using an optimized T7 exonuclease-based method

For endogenous tagging with long sequences using ssDNA donors, long ssDNA should be produced with high yield and purity. To this end, we optimized an ssDNA production method using dsDNA-specific T7 exonuclease for the preparation of long single-stranded HDR donors (Fig. 2a) (Nikiforov et al., 1994; Noteborn et al., 2021). First, dsDNA was amplified by PCR using HA-containing primers, whereas one of them has five sequential phosphorothioate (PS) bonds at the 5’ end. The amplified dsDNA was then column-purified and mixed with T7 exonuclease. Since the consecutive PS bonds block the digestion by T7 exonuclease, the strand with a non-modified 5’ end would be selectively digested, and the other strand would remain as an intact ssDNA. To verify that this strategy is effective enough to produce long ssDNA donors, dsDNA was amplified using different combinations of PS-modified (PS) and non-modified (noPS) primers and subsequently subjected to the T7 exonuclease reaction. The gel electrophoresis analysis showed that ssDNA bands were detected in the “PS-noPS” and “noPS-PS” conditions, where one of the primers was PS-modified, suggesting a successful production of ssDNA (Fig. 2b). However, we identified two major drawbacks of this method: first, dsDNA remained partially undigested even after the T7 exonuclease reaction. The second problem is that “PS-PS” dsDNA, in which both of the 5’ ends are PS-modified, seemed partially degraded by the exonuclease, suggesting that the protection by the PS modification was imperfect. We assumed that the latter problem of insufficient protection was due to the low efficiency of PS-modification of the long primers.

**Fig.2:**
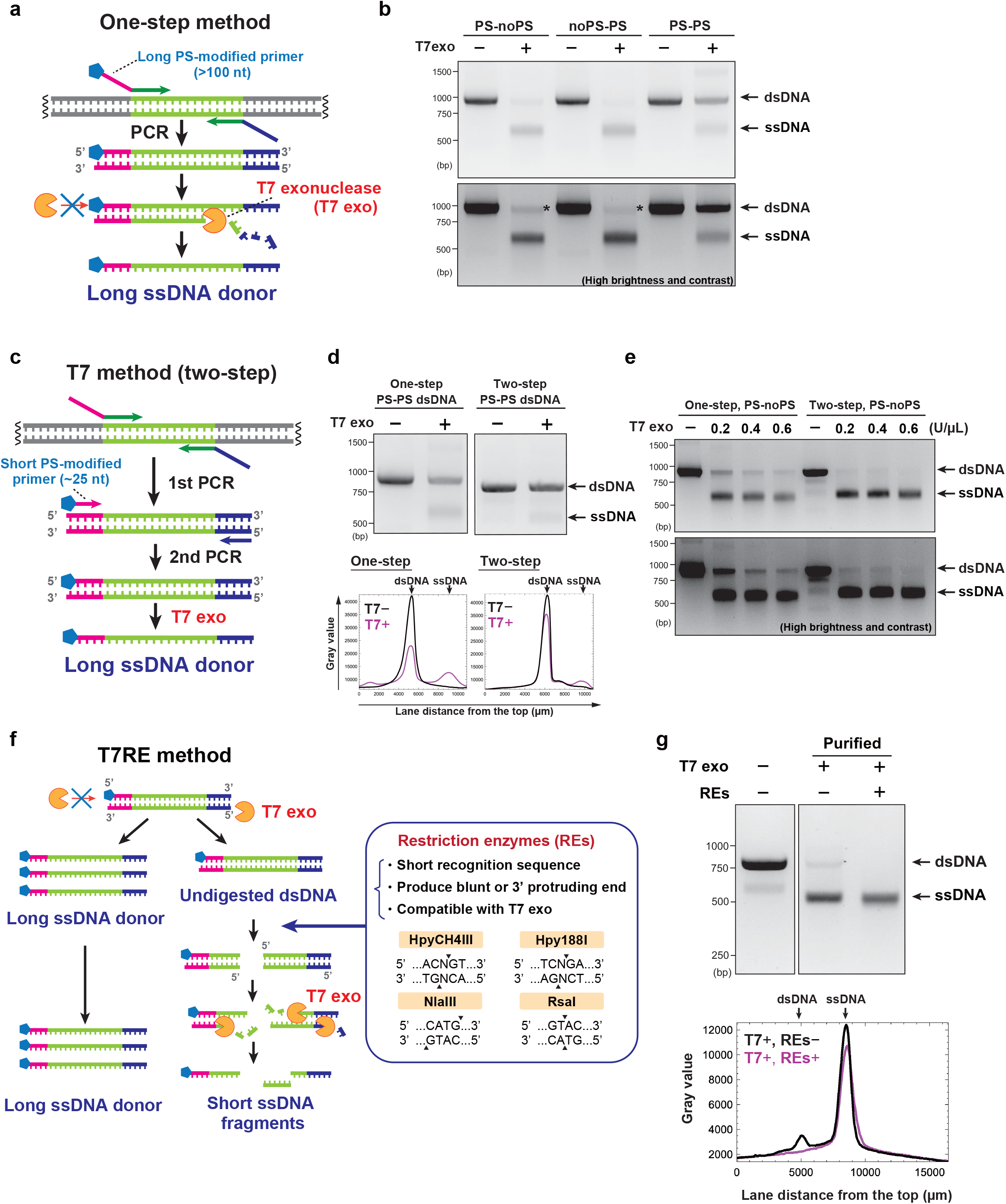
Optimization of long ssDNA production using T7 exonuclease and restriction enzymes. **a**, Schematic of long ssDNA production using T7 exonuclease (one-step method). dsDNA is amplified by PCR with a pair of long primers containing a 90 nt HA, one of which bears five consecutive PS bonds at the 5’ end. The strand with the non-modified 5’ end of the dsDNA is selectively digested by T7 exonuclease to produce long ssDNA donors. **b**, T7 exonuclease reaction on dsDNA amplified using three different combinations of primers (PS-modified (PS) or non-modified (noPS) for the forward and reverse primers). The bottom image is of the same gel as the top one, with higher brightness and contrast. ssDNA shows higher mobility than dsDNA of the same length. Asterisks show undigested dsDNA remnants. **c**, Schematic of ssDNA production by two-step PCR and T7 exonuclease (T7 method). The first PCR uses long non-modified primers to add the HAs, and the second uses a combination of short primers, one of which is PS-modified. **d**, “PS-PS” dsDNA was prepared with one-step or two-step PCR and subsequently subjected to the T7 exonuclease reaction. Plot profiles for each lane are shown below the gel electrophoresis images. **e**, Production of long ssDNA donors using the one-step and the two-step methods. The bottom image is of the same gel as the top one, with higher brightness and contrast. **f**, Schematic of ssDNA production using T7 exonuclease and restriction enzymes (T7RE method). After two-step PCR and T7 exonuclease reaction, the indicated four restriction enzymes digest dsDNA remnants to produce short dsDNA fragments which can be further degraded by T7 exonuclease. **g**, ssDNA production by the T7 and the T7RE methods. The last two lanes contain column-purified DNA products of both reactions. Plot profiles for the last two lanes are shown below the gel electrophoresis image.

To resolve this issue, we adopted a two-step PCR procedure to produce modified dsDNA, allowing the usage of short PS-modified primers, the purity of which is supposed to be higher than that of long primers (Fig. 2c). Compared to the initial one-step method, the degradation of the PS-PS dsDNA by T7 exonuclease was significantly reduced with the new two-step approach (Fig. 2d). When the two-step method was applied for long ssDNA preparation (i.e., with one modified 5’ end primer), the amount of undigested dsDNA was decreased, and the yield of ssDNA was higher (Fig. 2e). Furthermore, an annealing-based analysis revealed that the two-step method improved the strand selectivity of ssDNA production compared to the one-step method (Fig. S2).

Even though our improved T7 exonuclease-based method (hereafter referred to as T7 method) enables robust production of long ssDNA donors, a faint band of undigested dsDNA was still observed in agarose gel electrophoresis (Fig. 2e). For further improvement of the ssDNA purity, we added a restriction enzymes (REs) reaction after the digestion by T7 exonuclease (referred to as T7RE method) (Fig. 2f). We selected four REs (HpyCH4III, Hpy188I, NlaIII, and RsaI) whose recognition sequence is short so that they can be applied to various DNA sequences. Since these REs are fully active in the same buffer as T7 exonuclease, a sequential one-pot reaction can be applied to T7 exonuclease and REs. Importantly, all the four REs produce blunt or 3’-protruding ends, which serve as substrates for T7 exonuclease. Therefore, dsDNA fragments produced by REs are supposed to be degraded to even smaller ssDNA fragments in the presence of T7 exonuclease activity. Indeed, the T7RE method resulted in the successful removal of dsDNA remnants below the detection limit (Fig. 2g). In summary, our optimized T7RE method enables the preparation of long ssDNA donors with high yield and purity.

### Comparison of knock-in efficiency between dsDNA and ssDNA long donors

Using long ssDNA donors produced by the T7 and the T7RE methods (referred to as T7 donors and T7RE donors, respectively), we performed endogenous gene tagging in RPE1 cells (Fig. 3a). Electroporation of Cas12a-RNP and ssDNA donors for mNG tagging of HNRNPA1 or TOMM20 resulted in successful knock-in in each gene, as confirmed by the correct subcellular localization of the mNG signal (Fig. 3b). ssDNA donors prepared with our optimized methods were also applicable to Cas9-mediated mNG tagging of HNRNPA1 (Fig. 3b). We further performed flow cytometric analysis to evaluate the knock-in efficiency of ssDNA donors compared to that of dsDNA donors. For mNG tagging of HNRNPA1, knock-in efficiency tended to be lower with T7RE donors than with dsDNA donors, especially in the case of the sense strands (Fig. 3c, d). Similarly, for mNG tagging of TOMM20, the ratio of mNG positive cells introduced with T7RE donors was less than one-third of that with dsDNA donors (Fig. 3e). These data indicate low knock-in efficiency of T7RE donors compared to dsDNA donors across different target genes.

**Fig.3:**
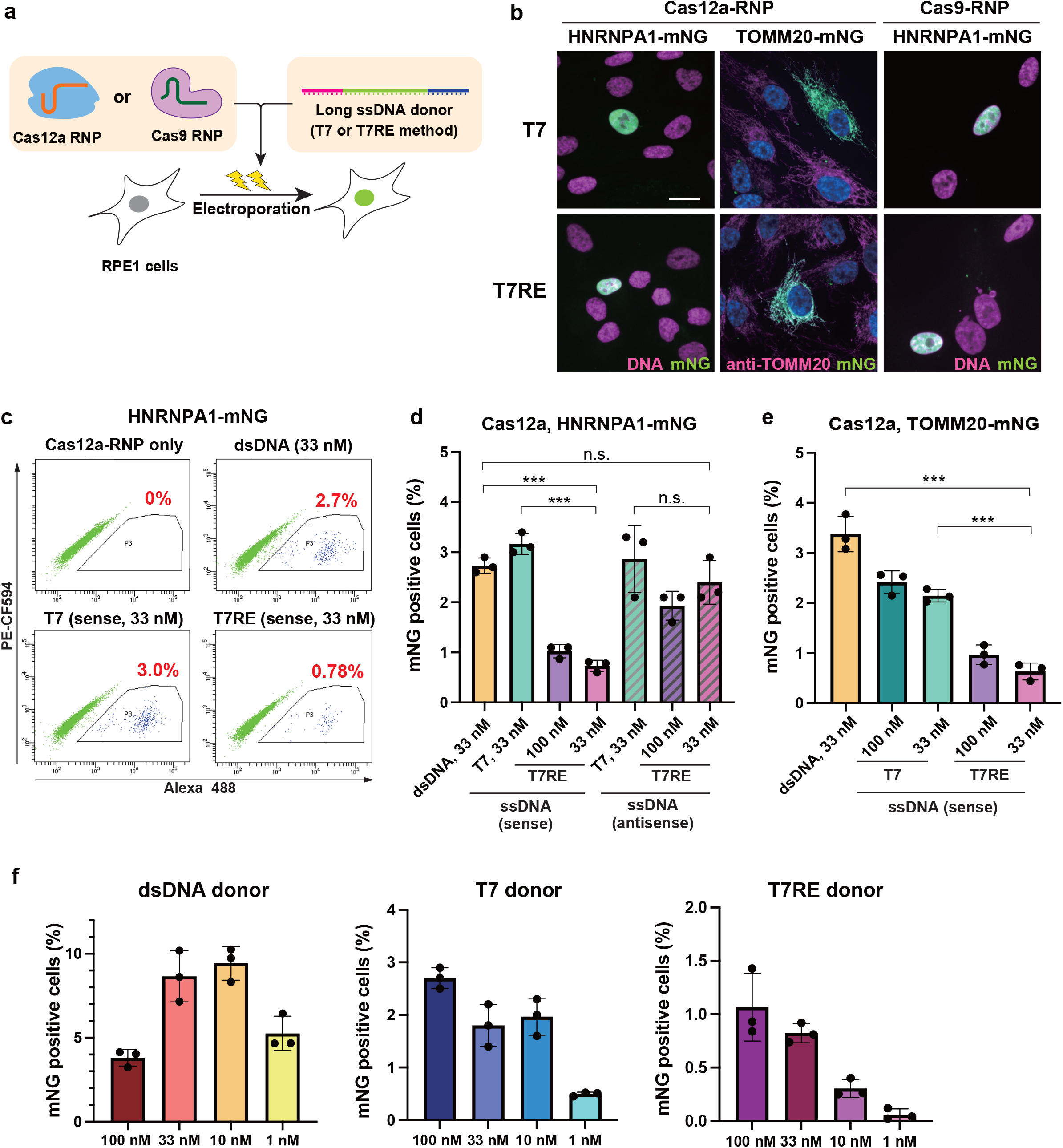
Comparison of knock-in efficiency between dsDNA and ssDNA long donors. **a**, Schematic of endogenous gene tagging using long ssDNA donors in RPE1 cells. ssDNA donors are produced by either the T7 or the T7RE method. **b**, Representative images of cells with mNG tagging to the indicated genes using T7 or T7RE donors. The indicated Cas nuclease was used for each condition. For ssDNA donors, sense strands were used. Cells at 7-13 days after electroporation were fixed and analyzed. Scale bar: 10 µm. **c**, Flow cytometric analysis of Cas12a-mediated HNRNPA1-mNG knock-in cells, using dsDNA, T7, and T7RE donors. Cells at 9 days after electroporation were analyzed. Percentages of cells with mNG signal are shown in the plots. **d**, Quantification of percentages of mNG positive cells from (**c**). Data from three biological replicates are shown. Approximately 10,000 cells were analyzed for each sample. **e**, Flow cytometric quantification of mNG positive cells in Cas12a-mediated TOMM20-mNG knock-in cells, using the indicated donors. Cells at 12 days after electroporation were analyzed. Data from three biological replicates are shown. >5,000 cells were analyzed for each sample. **f**, Titration of the indicated donors for mNG tagging of HNRNPA1 using Cas12a. Cells at 11 days (dsDNA) or 10 days (T7 and T7RE) after electroporation were analyzed. For ssDNA donors, sense strands were used. Data from three biological replicates are shown. Approximately 10,000 cells were analyzed for each sample. Data are presented as mean ± S.D. P-value was calculated by the Tukey–Kramer test. ***P <0.001, n.s.: Not significant.

Interestingly, the knock-in rate with T7RE donors tended to be also lower than that with T7 donors in all the tested conditions (Fig. 3c-e). The difference in efficiency between the two donors might be attributed to undigested dsDNA remnants in T7 donors. To estimate whether a small amount of dsDNA remnant would impact the knock-in performance of T7 donors, we conducted titration of donor concentration for dsDNA. Analysis by flow cytometry showed that dsDNA donors retained more than half of their maximum efficiency even at the concentration of 1 nM, while the efficiency of 1 nM of T7RE donors was reduced to less than one-tenth of their maximum (Fig. 3f). The result suggests that a small amount of dsDNA remnant among T7 donors might work as a template for HDR together with the ssDNA. Importantly, the concentration of dsDNA donors required to reach their maximum efficiency was lower than that of ssDNA. Collectively, these data indicate the superiority of dsDNA donors to ssDNA donors in terms of knock-in efficiency.

### Evaluation of the perfect HDR rate using long-read amplicon sequencing and *knock-knock* pipeline

Next, we compared the frequency of precise insertion of transgenes between dsDNA and ssDNA donors in RPE1 cells. To this end, we performed the long-read amplicon sequencing by PacBio and analysis by *knock-knock*, a computational framework that allows a high-throughput genotyping of knock-in alleles (Canaj et al., 2019). We applied this approach to mNG tagging of HNRNPA1 using Cas12a-RNP (Fig. 4a). After electroporation and subsequent cell expansion for two to three weeks, mNG positive cells were collected by fluorescence-activated cell sorting (FACS) to enrich knock-in cells. Genomic DNA was isolated after several days of culture in a 96-well plate. The specific DNA sequence at the target locus was then amplified for library preparation of sequencing. After sequencing by PacBio and Circular Consensus Sequence (CCS) generation, *knock-knock* classified each Hi-Fi read into a specific category of the knock-in outcome, such as WT, indels, HDR, or subtypes of mis-integration.

**Fig.4:**
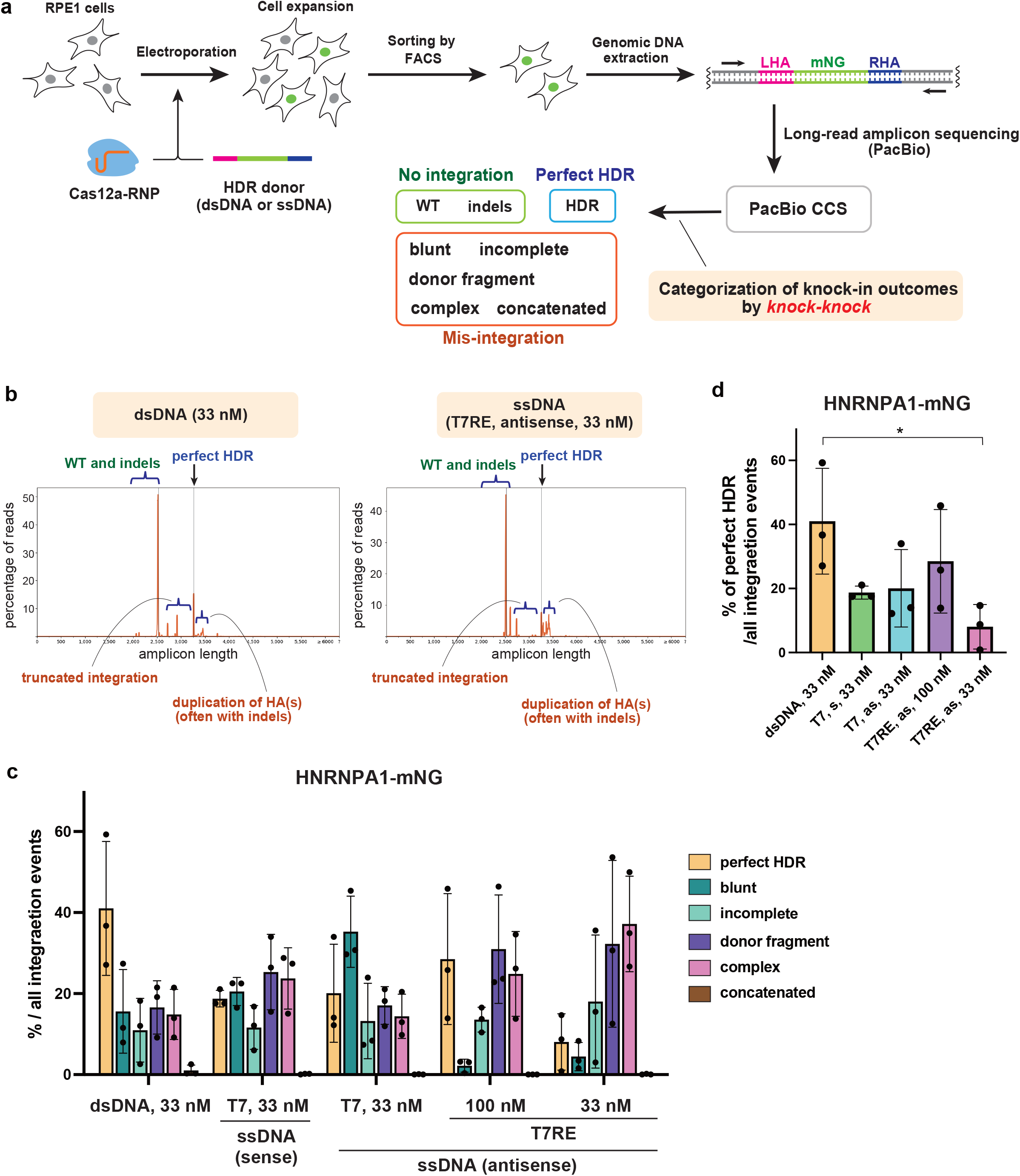
Long-read amplicon sequencing and *knock-knock* analysis to evaluate the accuracy of long transgene insertion with dsDNA and ssDNA donors. **a**, Schematic overview of analysis of knock-in outcomes. After electroporation of Cas12a-RNP and HDR donors for mNG tagging of HNRNPA1, cells were expanded for two to three weeks. mNG positive cells were then collected by FACS, and genomic DNA was isolated. Libraries for sequencing were prepared from the amplified target locus and subjected to long-read amplicon sequencing by PacBio. After analysis of sequencing outputs, including CCS generation, knock-knock categorized each read into a specific category of a knock-in outcome. **b**, Representative plots generated by *knock-knock* showing the distribution frequency of amplicon length. The range of read lengths corresponding to WT and indels, perfect HDR, truncated integrations, and duplication of homology arm(s) are indicated. **c**, Distribution of integration events across the donor types. For each category, the percentage within total integration events was calculated. Data from three biological replicates are shown. For each sample, 11559-44431 reads were categorized as the integration events from 43697-91850 total reads. **d**, Data from **c**, the frequencies of perfect HDR are highlighted. A two-tailed, unpaired Student’s t-test was used to obtain the P-value. *P < 0.05. s: sense strand, as: antisense strand.

The result of *knock-knock* analysis revealed that various mis-integration events occurred in addition to precise insertion via HDR for both dsDNA and ssDNA donors, as previously described (Canaj et al., 2019) (Fig. 4b, c). *Knock-knock* classified these mis-integration events into the following categories: blunt (at least one of the donor ends is directly ligated to the DSB site), incomplete (only one side of the donor is integrated via HDR), concatemer (multiple donors are inserted), donor fragment (both ends of the donor are integrated in a non-HDR manner), and complex (not classified into the other four mis-integration categories).

We then calculated the percentage of each category to total integration events. The results show that the rate of perfect HDR tends to be lower in ssDNA donors than in dsDNA donors (Fig. 4c, d). The blunt integration, which is assumed to be an outcome of NHEJ-based-ligation, was less likely to occur in T7RE (pure ssDNA) donors than in dsDNA donors (Fig. 4c). On the contrary, integration of donor fragments and complex integration were prominent in T7RE donors compared to dsDNA donors. Across the conditions, only a small percentage of reads were classified into the concatemer category. Taken together, these data obtained from *knock-knock* analysis suggest that dsDNA outperform ssDNA in the frequency of perfect HDR for long transgene insertion.

### Comparison of a propensity for homology-independent integration between dsDNA and ssDNA donors

The results of *knock-knock* analysis show that mis-integration of the mNG sequence to the target site is likely to occur more frequently than precise integration via perfect HDR regardless of the donor type, suggesting the prevalence of homology-independent integration. To further compare the propensity for homology-independent integration between dsDNA and ssDNA, we used donors without HAs for Cas12a-mediated mNG tagging of TOMM20 (Fig. 5a). When analyzed by flow cytometry, fluorescent cells were observed with T7RE donors at a similar level as dsDNA donors (Fig. 5b). Furthermore, mitochondrial localization of mNG fluorescence was confirmed by microscopic observation for all the conditions, suggesting integration of mNG donors into the targeted *TOMM20* locus via non-homologous pathways (Fig. 5c). It should be noted that the fraction of mNG positive cells is expected to be much lower than the frequency of total donor integration events at the target site because the mNG signal can be observed only when mNG is inserted in the correct orientation and a correct reading frame.

**Fig.5:**
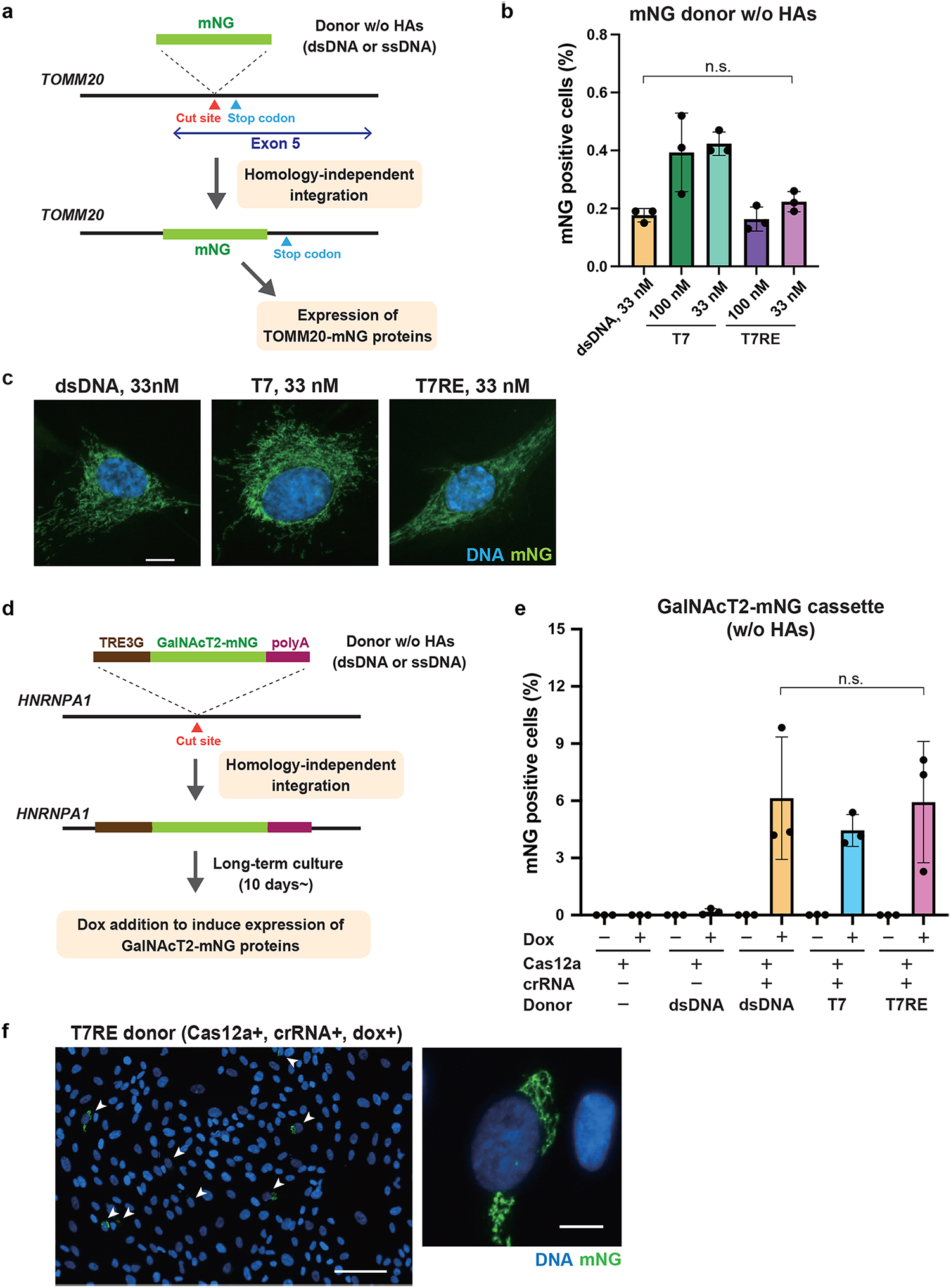
Comparison of homology-independent integration between dsDNA and ssDNA donors. **a**, Schematic for evaluating homology-independent integration of mNG donors into Cas nuclease-induced DSBs. Since the Cas12a cleavage site is located inside the coding region of the *TOMM20* gene, homology-independent integration of mNG into the cleavage site in the correct orientation and a correct reading frame leads to the expression of TOMM20-mNG proteins. **b**, Flow cytometric analysis of the homology-independent integration experiment. Sense strands were used for T7 and T7RE donors. Cells at 9 days after electroporation were analyzed. Data from three biological replicates are shown. >5,000 cells were analyzed for each sample. **c**, Representative images from the homology-independent integration experiment. Cells at 11 days after electroporation were fixed and analyzed. Scale bar: 10 µm. **d**, Schematic overview of the workflow for evaluating homology-independent integration of GalNAcT2-mNG cassettes into Cas nuclease-induced DSBs. Non-integrated cassettes are cleared from cells during long-term culture for more than 10 days. The cassettes inserted into the genome produce doxycycline (dox)-induced expression of a Golgi protein GalNAcT2-mNG. **e**, Flow cytometric analysis of the cassette integration experiment. Cells at 20 days after electroporation were treated with 1 µg/mL of doxycycline (dox) for 24h and analyzed. Data from three biological replicates are shown. Approximately 20,000 cells were analyzed for each sample. **f**, Representative images from the cassette integration experiment. Cells at 13 days after electroporation were treated with 1 µg/mL doxycycline (dox) for 24h and fixed for analysis. Arrowheads indicate cells with Golgi-like mNG signals. Scale bar: 100 µm (the left panel), 10 µm (the right panel). Data are presented as mean ± S.D. A two-tailed, unpaired Student’s t-test was used to obtain the P-value. n.s.: Not significant.

To further confirm the occurrence of homology-independent integration, we prepared a DNA cassette encoding TRE3G-GalNAcT2-mNG-polyA, which allows doxycycline-dependent expression of mNG-fused Golgi protein GalNAcT2 (Fig. 5d). This approach was expected to be more sensitive than the former strategy (Fig. 5a) for detecting integration events because GalNAcT2-mNG expression occurs regardless of how the cassette is inserted into the genome due to its own promoter. The electroporated cells were cultured for more than 10 days so that the non-integrated cassettes could be cleared from the cells. When analyzed by flow cytometry, the control population that was electroporated with Cas12a and dsDNA cassettes but without crRNA exhibited about 0.2% of fluorescent cells, presumably due to random integration of dsDNA cassettes into the genome, as reported previously (Fig. 5e) (Fueller et al., 2020). Nevertheless, a higher population of mNG positive cells (4 to 10%) was detected from the cells electroporated with Cas12a-crRNA RNP complexes and dsDNA cassettes, suggesting the integration into the Cas12a-induced DSBs. In the case of ssDNA donors, the fractions of mNG positive cells were comparable to those of dsDNA, consistent with the result of the former analysis (Fig. 5b). The mNG signals detected by flow cytometry were confirmed to be derived from the GalNAcT2-mNG cassette since the fluorescence was doxycycline-dependent and a Golgi-like localization of the mNG signal was observed by microscopy (Fig. 5f). Therefore, it is likely that there are no significant differences between dsDNA and ssDNA donors in their propensity for homology-independent insertion into the Cas nuclease-induced DSBs. In conclusion, our comprehensive analyses indicate that ssDNA donors are not superior to dsDNA for endogenous gene tagging with long transgenes in RPE1 cells.

## Discussion

In this study, we systematically compared the performance of dsDNA and ssDNA donors for CRISPR-Cas knock-in of long transgenes in human diploid RPE1 cells. Our analysis revealed that knock-in efficiency tended to be higher for dsDNA compared to the pure ssDNA (T7RE) donors in RPE1 cells. Recent studies have shown that long ssDNA donors can be used for efficient knock-in in various species (Li et al., 2017; Miura et al., 2015; Nakayama et al., 2020; Quadros et al., 2017). Especially in zebrafish, ssDNA has been shown to be more efficient than dsDNA as long HDR donors (Ranawakage et al., 2021). On the other hand, in human cells such as primary T cells, HEK293T cells, and iPS cells, the knock-in efficiency of long ssDNA donors has been described to be lower than that of dsDNA donors (Canaj et al., 2019; Li et al., 2017; Roth et al., 2018). Thus, our data together with these previous reports indicate that dsDNA outcompetes ssDNA for knock-in efficiency in human cells.

To establish cell lines with accurate gene knock-in, the efficiency of perfect HDR is crucial rather than seeming knock-in efficiency just assessed by flow cytometry or microscopy. By performing long-read amplicon sequencing and *knock-knock* analysis, we quantified the frequency of precise insertion via perfect HDR among a pool of heterogeneous repair outcomes in a high throughput manner, as described previously (Canaj et al., 2019). Our data show that long ssDNA donors result in lower percentages of perfect HDR in RPE1 cells than dsDNA donors. This observation is consistent with the previous study by Canaj and colleagues, who developed *knock-knock*, in which the perfect HDR rate for long ssDNA donors was similar to or lower than dsDNA donors in three different cell lines. Therefore, dsDNA donors are presumably superior to ssDNA donors in terms of precise knock-in of long transgenes.

The previous *knock-knock* data show that dsDNA donors are more prone to NHEJ-mediated mis-integration into the target locus (Canaj et al., 2019). In agreement with this, our data from *knock-knock* analysis showed that the percentage of the blunt integration of dsDNA donors was higher than that of T7RE donors (pure ssDNA). On the other hand, the previous study reported that ssDNA donors show more pronounced incomplete mis-integration, in which one end exhibits HDR and the other is repaired imperfectly (often in a truncated manner), which however could not be confirmed by our analysis. This difference might arise due to our enrichment procedure of fluorescent cells by FACS prior to the sequencing, which eliminates cells with truncated integrations that did not express functional fluorescent proteins.

Linear dsDNA is prone to be randomly integrated into the genome via non-HDR pathways at sites of naturally occurring DSBs (Saito et al., 2017; Zelensky et al., 2017). In the context of endogenous tagging using long HDR donors, previous reports suggest that non-homologous integration or off-target integration is less likely to occur with ssDNA than linear dsDNA (Li et al., 2017; Roth et al., 2018). However, our two different analyses of homology-independent integration revealed that the integration rate of donors without HAs is almost the same for ssDNA and dsDNA, suggesting comparable propensity for off-target integration of these donors. Consistently, our *knock-knock* data showed that the frequency of imprecise integration of ssDNA donors was not lower than that of dsDNA. Thus, considering the knock-in efficiency, the insertion accuracy and the off-target integration frequency, ssDNA is probably not superior to dsDNA as long HDR donors for knock-in in human cell lines. Given that dsDNA donors can be prepared easier than long ssDNA donors, we suggest using dsDNA rather than ssDNA as HDR donors for endogenous tagging with long transgenes in human cells.

## Acknowledgments

We thank Miho Kiyooka and Wei Chen at National Institute of Genetics for supporting NGS sequencing, Dr. Yusuke Kishi at Institute for Quantitative Biosciences at the University of Tokyo for supporting quality control of NGS library preparation, and the Kitagawa lab members and Dr. Elmar Schiebel from ZMBH in Heidelberg University for technical supports and helpful discussions. This work was supported by JSPS KAKENHI grants (Grant numbers: 18K06246, 19H05651, 20K15987, 20K22701, 21H02623, 22H02629) from the Ministry of Education, Science, Sports and Culture of Japan, the PRESTO program (JPMJPR21EC) of the Japan Science and Technology Agency, Takeda Science Foundation, The Uehara Memorial Foundation, The Research Foundation for Pharmaceutical Sciences, Koyanagi Zaidan, The Kanae Foundation for the Promotion of Medical Science, Kato Memorial Bioscience Foundation, and Tokyo Foundation for Pharmaceutical Sciences.

## Author contributions

S.H. conceived and designed the study. A.M. designed and performed most of the experiments. M.G. optimized the genome editing conditions. C.T. performed tagging of CAMSAP2 and validated knock-in specificity. K.K.I. and A.M. analyzed the PacBio data with *knock-knock*. M.H. performed endogenous tagging of p53. T.K. contributed to quantification by flow cytometry. M.F. and T.C. provided suggestions. A.T. performed PacBio sequencing. A.M., S.H. and D.K. analyzed the data. A.M., S.H. and D.K. wrote the manuscript. All authors contributed to discussions and manuscript preparation.

## Competing financial interests

The authors declare no competing financial interests.

## Materials and methods

### Cell culture

RPE1 cells were cultured in Dulbecco’s Modified Eagle Medium/Nutrient Mixture F-12 (DMEM/F-12) supplemented with 10% FBS, 100 U/mL penicillin, and 100 µg/mL streptomycin at 37°C in a humidified 5% CO_2_ incubator.

### dsDNA donor preparation

dsDNA donors were amplified by PCR from a plasmid encoding the 5xGA linker-mNG sequence using two primers containing 90-base left and right HA sequences, respectively. We used Q5 High-Fidelity 2X Master Mix (New England Biolabs) for PCR. DpnI (0.04 U/µL) and Exonuclease I (0.4 U/µL) purchased from New England Biolabs were directly added to the PCR reaction mix and incubated at 37°C for 30 min, followed by heat inactivation at 80°C for 20 min. DpnI and Exonuclease I were used for digestion of residual template plasmids and primers, respectively. The dsDNA donors were then column-purified using the NucleoSpin Gel and PCR Clean-up kit (Macherey-Nagel) and stored at -20°C or directly used for electroporation. All primer sequences used in this study are listed in Supplementary Table1.

### ssDNA production using one-step PCR and T7 exonuclease

dsDNA was amplified by PCR as described above with a minor alteration. One of the HA-containing primers was modified with five consecutive phosphorothioate (PS) bonds at the 5’ end. The dsDNA was treated with DpnI and Exonuclease I and column-purified. T7 exonuclease (0.3 U/µL, New England Biolabs) was mixed with the purified dsDNA (60 ng/µL) in rCutSmart buffer and incubated at 25°C for 30 min. The reaction mix was directly used for electrophoresis using 2% agarose gel supplemented with Midori Green Advance (Nippon Genetics) at 4°C for 40 to 60 min to check ssDNA production.

### ssDNA donor preparation with the T7 or T7RE method

dsDNA was prepared with two-step PCR. The first-round PCR was performed using non-modified primers followed by treatment with DpnI and Exonuclease I, as described above. The specific PCR product was gel purified and then used as a template for the second-round PCR with two short primers (about 25 nt), one of which contains five sequential PS bonds at the 5’ end. After column purification, the dsDNA was reacted with T7 exonuclease as described above. For the preparation of T7RE donors, HpyCH4III (0.025 U/µL), Hpy188I (0.1 U/µL), NlaIII (0.025 U/µL), and RsaI (0.05 U/µL) purchased from New England Biolabs were added directly to the T7 exonuclease reaction mix and incubated at 37°C for 15 min. After the enzymatic reactions, ssDNA was column-purified using Buffer NTC (Macherey-Nagel) as a binding buffer. Typically, 4 to 5 µg of ssDNA was obtained from 15 µg of dsDNA, when elution with 15 µL of nuclease-free water was conducted twice.

### Synthesis of guide RNA

Guide RNA (sgRNA for Cas9 and crRNA for Cas12a) was transcribed in *vitro* from PCR-generated DNA templates according to a previous method (Komori et al., 2021). Briefly, for sgRNA, template DNA containing T7 promoter and sgRNA sequence was amplified by PCR from five different oligos. Template DNA for crRNA was likewise assembled by PCR from two different oligos. The purified DNA template was subjected to *in vitro* transcription by T7 RNA polymerase using the HiScribe T7 High Yield RNA Synthesis Kit (New England Biolabs). After being treated with DNase I (Takara Bio), the synthesized guide RNA was purified using the RNA Clean & Concentrator Kit (Zymo Research). All guide RNA sequences used in this study are listed in Supplementary Table2.

### Gene knock-in using CRISPR-Cas12a and CRISPR-Cas9 system

Endogenous gene tagging using the CRISPR-Cas12a system was performed with the electroporation of Cas12a-RNP and HDR donors (dsDNA, T7, or T7RE donors) using the Neon Transfection System (Thermo Fisher Scientific) according to the manufacturer’s protocol. A.s. Cas12a Ultra (1 µM) from Integrated DNA Technologies (IDT) and crRNA (1 µM) were pre-incubated in resuspension buffer R (Thermo Fisher Scientific) at room temperature and mixed with cells (0.125 ×10^5^ /µL), Cpf1 electroporation enhancer (1.8 µM, IDT), and the HDR donors (33 nM). Electroporation was conducted using a 10 µL Neon tip at a voltage of 1300 V with two 20 ms pulses. The transfected cells were seeded into a 24-well plate.

CRISPR-Cas9-mediated knock-in was performed similarly to the Cas12a-RNP condition described above, with a modification in the electroporation solution. Briefly, HiFi Cas9 protein (1.55 µM, IDT) and sgRNA (1.84 µM) were pre-incubated in buffer R and mixed with cells, Cas9 electroporation enhancer (1.8 µM, IDT), and the HDR donors.

### Quantification of knock-in efficiency by flow cytometry

Flow cytometric analysis was conducted 5 to 12 days after electroporation. Cells were harvested with trypsin/EDTA solution and suspended in DMEM/F12 medium with HEPES and without phenol red. The cell suspensions were analyzed using BD FACS Aria III (BD Biosciences), equipped with 355/405/488/561/633 nm lasers to detect cells with mNG signal. Data were collected from more than 5,000 gated events.

### Amplicon sequencing and analysis by knock-knock

#### Genomic DNA preparation

After electroporation of Cas12a-RNP targeting the *HNRNPA1* locus and HDR donors, cells were expanded for 17 days. mNG positive cells were sorted using BD FACS Aria III and seeded into a 96-well plate. Cells were expanded until confluent and genomic DNA was extracted using DNAzol Direct (Molecular Research Center).

#### Amplicon sequencing

Amplicon libraries were prepared with two-step PCR and subsequent adapter ligation, according to the protocol provided by Pacific Biosciences (Part Number 101-791-800 Version 02 (April 2020)) with slight modifications. The first-round PCR was conducted to amplify a region flanking the target site of mNG insertion from extracted genome DNA. For the amplification, KOD One Master Mix (TOYOBO) was used with primers tailed with universal sequences which serve as an annealing site for a barcoded primer. The amplified DNA was purified using AMPure XP (Beckman Coulter). The purified DNA was re-amplified by PCR using primers from Barcoded Universal F/R Primers Plate-96v2 (Pacific Biosciences) and subsequently purified with AMPure PB beads (Pacific Biosciences). The barcoded amplicons were then analyzed by TapeStation (Agilent Technologies) and Qubit Fluorometer (Thermo Fisher Scientific). All the amplicons were pooled as one sample in equimolar amounts. A pooled sequencing library was prepared using the SMRTbell Express Template Prep Kit 2.0 (Pacific Biosciences). One Sequel II SMRT cell was run on the PacBio Sequel II Platform with Binding Kit 2.0/Sequencing Kit 2.0 and 24 hr movies, yielding a total of 7,031,124 polymerase reads (328,085,638,677 bp). The consensus reads (1,572,695 HiFi reads with QV ≧40) were generated from the raw full-pass subreads using the PacBio CCS program (SMRT Link v10.2.1.143962) and then 1,319,631 barcoded reads were selected after demultiplexing.

#### Analysis of knock-in outcomes by knock-knock

Before analysis of knock-in outcomes, the universal primer sequences at both ends were trimmed from the reads. We then analyzed these trimmed reads with *knock-knock*, a computational pipeline developed by Canaj et al. (2019). The source code is available at https://github.com/jeffhussmann/knock-knock.

### Analysis of homology-independent integration using GalNAcT2-mNG cassettes

The TRE3G-GalNAcT2 (1-114 aa)-5xGA-mNG-BGH polyA sequence was amplified by PCR from pRetroX-TRE3G-GalNAcT2-mNG-polyA plasmid for the preparation of donor cassettes, using primers not having HA sequences. The PCR products were subjected to the preparation of dsDNA and ssDNA donors as described above. The purified DNA cassettes (33nM) were electroporated into RPE1-Tet3G cells with Cas12a-RNP targeting the *HNRNPA1* locus. Electroporated cells were cultured for more than 10 days to remove the non-integrated cassettes. The cells at 13 days and 20 days after electroporation were treated with doxycycline (1 µg/mL, Merck) for 24 hours and subjected to microscopic and flow cytometric analyses, respectively.

### Genomic PCR

Genomic DNA was purified using the NucleoSpin DNA RapidLyse kit (Macherey-Nagel). The knock-in region was amplified by PCR using primers and KOD One PCR Master Mix, and then subjected to agarose gel electrophoresis.

### Immunofluorescence

For indirect immunofluorescence, cells cultured on coverslips (Matsunami Glass) were fixed with 4% PFA in PBS at room temperature for 15 min. Fixed cells were blocked with blocking buffer (1% bovine serum albumin in PBS containing 0.05% Triton X-100) for 30 min at room temperature. The cells were then incubated with primary antibodies in the blocking buffer at room temperature for 1 hour in a humid chamber. After washing with PBS, the cells were incubated with secondary antibodies in the blocking buffer at room temperature for 30 min. The coverslips were washed with PBS and mounted onto glass slides (Matsunami Glass) using ProLong Gold Antifade Mountant with DAPI (Thermo Fisher Scientific), with the cell side down.

### Western blotting

Cells were lysed on ice in lysis buffer containing 50 mM Tris-HCl, pH 8.0, 150 mM NaCl, 1% NP-40, 0.5% Sodium deoxycholate, 0.1% SDS, 5 mM EDTA,15 mM MgCl_2_, 1:1,000 protease inhibitor cocktail (Nacalai Tesque), and 1:1,000 phosphatase inhibitor cocktail (Nacalai Tesque). After centrifugation, the supernatant mixed with Laemmli sample buffer was boiled and subjected to SDS-PAGE. Separated proteins were transferred onto Immobilon-P PVDF membrane (Merck) using Trans-Blot SD Semi-Dry Electrophoretic Transfer Cell (Bio-Rad Laboratories). The membrane was blocked with 2% skim milk in PBS containing 0.02% Tween-20 and probed with the primary antibodies, followed by incubation with their respective HRP-conjugated secondary antibodies. The membrane was soaked with Chemi-Lumi One L or Chemi-Lumi One Super (Nacalai Tesque) for the signal detection using ChemiDoc XRS+ (Bio-Rad Laboratories).

### Antibodies

The following primary antibodies were used in this study: anti-TOMM20 (Santa Cruz Biotechnology; sc-17764, IF 1:1,000), anti-mNG (Chromotek, 32f6; IF 1:500), anti-GM130 (Cell Signaling Technology, #12480; IF 1:1,000), anti-HNRNPA1 (Santa Cruz Biotechnology, sc-32301; WB 1:500), anti-mNG (Cell Signaling Technology, #53061, WB 1:100), and anti-HSP90 (BD Biosciences, 610419; WB 1:5,000). The following secondary antibodies were used: donkey anti-mouse IgG Alexa Fluor 555 (Invitrogen, A32773; IF 1:500), donkey anti-mouse IgG Alexa Fluor 647 (Invitrogen, A32787; IF 1:500), donkey anti-rabbit IgG Alexa Fluor 647 (Invitrogen, A32795; IF 1:500), anti-mouse IgG HRP (Promega, WB 1:10,000), and anti-rabbit IgG HRP (Promega, WB 1:10,000)

### Statistical analysis

Statistical comparison between the data from different groups was performed in PRISM v.9 software (GraphPad) using either the Tukey–Kramer test or a two-tailed, unpaired Student’s t-test as indicated in the figure legends. P-value <0.05 was considered statistically significant. All data shown are mean ± S.D. Sample sizes are indicated in the figure legends.

## Figure legends

**Supplementary Table 1:**
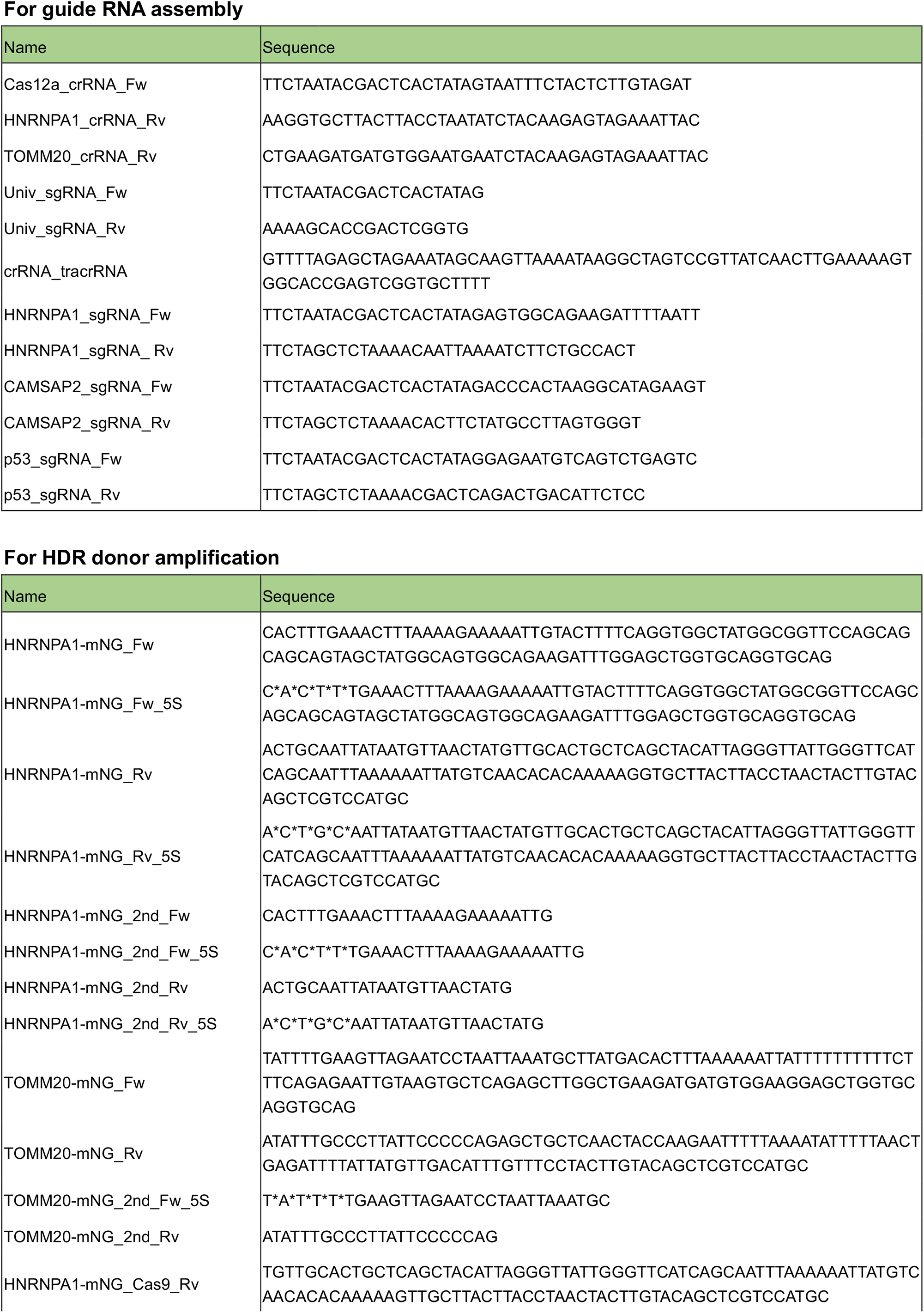
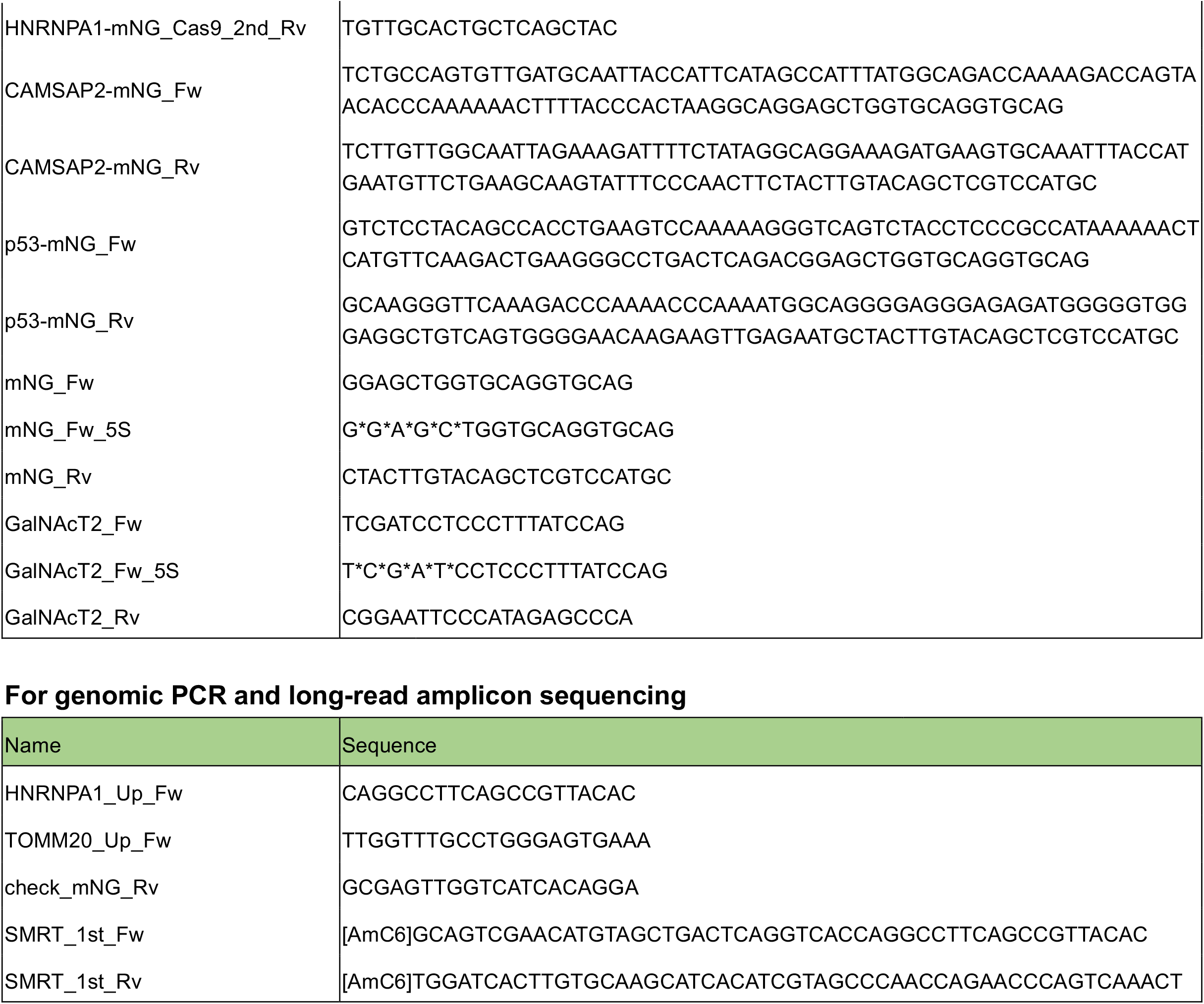
Primer sequences for PCR.

**Supplementary Table 2:**
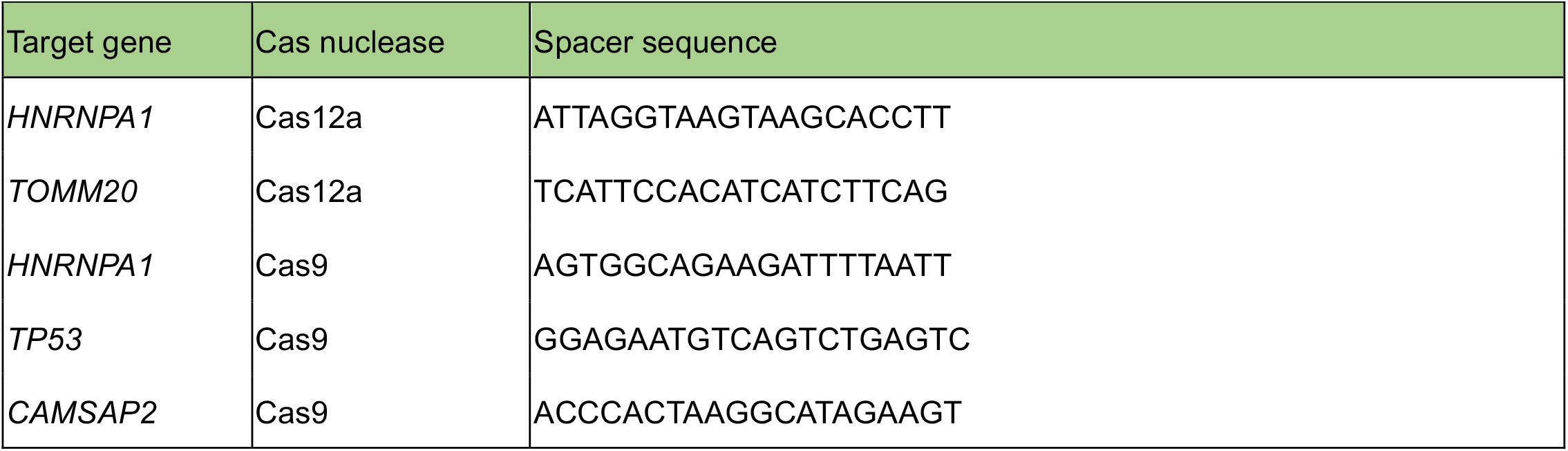
Guide RNA sequences.

## Notes

### Competing Interest Statement

The authors have declared no competing interest.

